# Characterization in mice of the stromal niche maintaining AT2 stem cell self-renewal in homeostasis and disease

**DOI:** 10.1101/2021.01.28.428090

**Authors:** Sara Taghizadeh, Monika Heiner, Jochen Wilhelm, Susane Herold, Chengshui Chen, JinSan Zhang, Saverio Bellusci

## Abstract

Resident mesenchymal cells (rMCs defined as Cd31^Neg^Cd45^Neg^Epcam^Neg^) control the self-renewal and differentiation of alveolar epithelial type 2 (AT2) stem cells in vitro. The identity of these rMCs is still elusive. Among them, Axin2^Pos^ mesenchymal alveolar niche cells (MANCs), which are expressing Fgf7, have been previously described. We propose that an additional population of rMCs, expressing Fgf10 (called rMC-Sca1^Pos^Fgf10^Pos^) are equally important to maintain AT2 stem cell self-renewal.

The alveolosphere model, based on the AT2-rMC co-culture in growth factor reduced Matrigel, was used to test the efficiency of different rMC subpopulations isolated by FACS from adult murine lung to sustain the self-renewal and differentiation of AT2 stem cells.

We demonstrate that rMC-Sca1^Pos^Fgf10^Pos^ cells are efficient to promote the self-renewal and differentiation of AT2 stem cells. Co-staining of adult lung for *Fgf10* mRNA and Sftpc protein respectively, indicate that 28% of Fgf10^Pos^ cells are located close to AT2 cells. Co-ISH for *Fgf7* and *Fgf10* indicate that these two populations do not significantly overlap. Gene arrays comparing rMC-Sca1^Pos^Axin2^Pos^ and rMC-Sca1^Pos^Fgf10^Pos^ support that these two cell subsets express differential markers. In addition, rMC function is decreased in diabetic and obese *ob/ob* mutant compared to WT mice with a much stronger loss of function in males compared to females.

In conclusion, rMC-Sca1^Pos^Fgf10^Pos^ cells play important role in supporting AT2 stem cells self-renewal and differentiation. This result sheds a new light on the subpopulations of rMCs contributing to the AT2 stem cell niche in homeostasis and in the context of COVID-19 pathogenesis.

**Key message:** *What is already known about the subject?:* Resident mesenchymal cells (rMCs defined as Cd31^Neg^Cd45^Neg^Epcam^Neg^) control the self-renewal and differentiation of alveolar epithelial type 2 (AT2) stem cells in vitro. The identity of these rMCs is still elusive. Among them, Axin2^Pos^ mesenchymal alveolar niche cells (MANCs), which are expressing Fgf7, have been previously described.

*What does this study add?:* Our study shows that an additional population of rMCs, expressing Fgf10 (called rMC-Sca1^Pos^Fgf10^Pos^) is equally important to maintain AT2 stem cell self-renewal. rMC-Sca1^Pos^Fgf10^Pos^ are LipidTox^High^ and are located close to AT2s. In addition, rMC-Sca1^Pos^Fgf10^Pos^ cells support AT2 stem cell self-renewal and differentiation thereby identifying these cells as *bone fide* functional lipofibroblasts (LIFs). We have previously reported that LIF can transdifferentiate into activated MYF in the context of bleomycin-induced fibrosis in mice [1] and that activated MYF isolated from the lungs of end stage idiopathic fibrosis human patients can respond to Metformin to undergo transdifferentiation back to the LIF phenotype [2]. We also show that the function of rMCs-Sca1^Pos^ is negatively impacted by gender and obesity, which represent two major aggravating factors for COVID-19 pathogenesis, leading to either death or major complications after infection recovery such as lung fibrosis.

*How might this impact on clinical practice and future development?:* By establishing that rMC-Sca1^Pos^Fgf10^Pos^ are different from the MANCs, our study opens the way for a new key mesenchymal cell population that should be targeted to either prevent or reverse fibrosis. In addition, as this population maintains the AT2 stem cells self-renewal and differentiation, such targeting will also allow to progressively recover the loss in respiratory function.

## Introduction

Lung fibrosis is characterized by an accumulation in the respiratory airways of mesenchymal cells, called activated myofibroblasts (MYF) which over time, lead to impaired lung function [3]. Through the use of single cell transcriptomic approach, the heterogeneity of the different resident mesenchymal (rMC) populations present in the lung is starting to emerge. A key stromal cell is represented by the lipofibroblasts (LIFs), which are rich in lipid-droplets and can be stained and isolated using the vital dye LipidTox (LT). They express *Perilipin 2, platelet derived growth factor receptor alpha* (*Pdgfra*) and are negative for Acta2 and most importantly are located close to alveolar type 2 cells (AT2) [4]. They are proposed to supply AT2s with the triglycerides needed for the elaboration of surfactant. Using a co-culture assay of Pdgfrα^High^ mesenchymal cells and AT2s in growth factor reduced Matrigel, it has been proposed that the rMC-LIFs are essential for the maintenance of the self-renewal and differentiation of AT2 stem cells [5].

Fibroblast growth factor 10 (Fgf10) is a mesenchymal-specific gene expressed in the distal part of the embryonic lung during the early pseudoglandular stage [6]. Our lineage tracing analyses have shown that a subpopulation of Fgf10^Pos^ cells at embryonic day 12.5 serves as progenitor for rMC-LIFs at later stages of lung development. Additionally, in postnatal lungs, around 30% of rMC-LIF express the *Fgf10* gene [7]. The relevance of these rMC-Fgf10^Pos^ LIF in maintaining the self-renewal and differentiation of AT2 stem cells remains to be demonstrated. Interestingly, another population of stromal cells, called MANC (Mesenchymal Alveolar Niche Cells), which are positive for *Axin2, Pdgfra, Wnt2, Il6* and *Fgf7* [8], were also described to locate close to AT2 cells. How different are the rMC-Fgf10^pos^ LIFs to the MANC is still unclear.

Using lineage tracing, we previously reported the reversible transdifferentiation of the LIFs into activated MYF during fibrosis formation and resolution [1]. In addition, Metformin, a first line antidiabetic drug has been reported to enhance the activated MYF to LIF transition both in vitro and in vivo [2]. Repetitive damages to AT2s have been proposed to be linked to the development of lung fibrosis. Interestingly, one of the major and long-term complication for patients who survived SARS-CoV-2 infection (leading to COVID-19 disease) is fibrosis [9,10]. SARS-CoV-2 binds to its receptor angiotensin converting enzyme (ACE) mostly abundantly expressed in the AT2s. Massive damages to the AT2s likely unleashes the associated LIFs to become activated MYFs, thereby leading to fibrosis. Pertinent to this background, obesity/diabetes and gender have been proposed to be predictive factors for the severity of the disease [11,12].

In this paper, we used the in vitro alveolosphere model based on the AT2-rMC co-culture in growth factor reduced Matrigel to test the efficiency of different rMC subpopulations isolated by FACS from adult murine lung. We used antibodies against Cd45, Cd31, Epcam and Stem cell antigen 1 (Sca1) to refine the initial rMC population responsible for the maintenance of AT2 stem cells. Using a fluorescent substrate for ß-galactosidase activity, rMC-Sca1^Pos^Fgf10^Pos^ as well as rMC-Sca1^Pos^Axin2^Pos^ cells were sorted. Additional sorting was achieved using LT staining for cells containing high level of neutral lipids. After 2 weeks of co-culture, organoid size and colony forming efficiency were quantified. RNAscope with *Fgf7*- and *Fgf10*-labelled riboprobes, to label rMC-Axin2^Pos^ and rMC-Fgf10^Pos^ respectively, combined with Sftpc immunofluorescence on adult lungs was carried out. Gene array analyses were used to characterize sorted rMC-Sca1^Pos^Fgf10^Pos^ vs. rMC-Sca1^Pos^Axin2^Pos^. We used males and females C57BL6 and corresponding *ob/ob* mutant mice to determine the impact of obesity and gender on the functional capacity of the corresponding rMCs-Sca1^Pos^ to sustain AT2 stem cells.

Our results indicate that an rMC-Sca1^Pos^Fgf10^Pos^/LIF^Pos^ subpopulation is essential for the self-renewal of the AT2 stem cells. This subpopulation is likely different from the previously described MANC. In addition, obesity and gender impact the capacity of rMCs to maintain the self-renewal of AT2 stem cells indicating that future therapeutic approaches should be focussed on restoring the function of the mesenchymal niche.

## Materials and Methods

### Mice

*Sftpc*^*CreERT2/+*^ knock in (gift from Harold Chapman, UCSF), *tdTomato*^*flox*^ reporter (Stock 007908, Jacksonlab), *Fgf10*^*LacZ*^ reporter (Mailleux et al., 2005), *Axin2*^*LacZ*^ reporter (stock 009120, Jackson lab), *Lep*^*ob/ob*^ (aka *ob/ob*) mutant (stock 000632, Jackson lab) and wild type mice were maintained on the C57BL/6 background. All animal studies were performed according to protocols approved by the Animal Ethics Committee of the Regierungspraesidium Giessen (permit numbers: G7/2017–No.844-GP and G11/2019–No. 931-GP).

### Lung dissociation and Fluorescence-Activated Cell Sorting

See supplementary methods

### LipidTOX™ staining

See supplementary methods

### FluoReporter™ *lacZ* Flow Cytometry

Fluorescein di (b-D-galactopyranoside) (aka FDG) (Thermo Fischer Scientific #F1930) was used to isolate by FACS, cells expressing ß-galactosidase from *Fgf10*^*LacZ*^ and *Axin2*^*LacZ*^ reporter lines. Lungs were collected from 6-8 weeks old *Fgf10*^*LacZ*^ and *Axin2*^*LacZ*^ reporter mice. According to manufacturer’s instruction, single-cell suspension and FDG working solution were prewarmed and the cells were resuspended with chloroquine followed by loading by FDG. After incubation for 20 minutes, FDG loading is stopped by adding ice cold staining medium containing propidium iodide and chloroquine. Cells are then placed on ice and incubated with antibodies against Cd45, Cd31, Sca1, and Epcam (for details see Lung dissociation and Fluorescence-Activated Cell Sorting) before sorting using the FACSAria™ III (BD Bioscience) cell sorter. Cells were sorted through a flow chamber with a 100-μm nozzle tip under 25 psi sheath fluid pressure. Cells were collected in sorting media (advanced DMEM:F12 (gibco#12634-010) plus 10% FBS and 1% P/S).

### Alveolar Organoid Assay

5000 Lyso^Pos^Tom^Pos^ cells (AT2s from adult *Sftpc*^*CreERT2/+*^; *tdTomato*^*flox/+*^ lungs) and 50000 rMCs cells were resuspended in 100 µl culture medium (sorting media plus 1% ITS (gibco # 41400-045)) and mixed 1:1 with 100 µl growth factor-reduced phenol Red-free Matrigel (Corning #356231). Cells were seeded in individual 24-well 0.4 µm Transwell inserts (Falcon, SARSTEDT). After incubation at 37°C for 15 minutes, 500 µl of culture was placed in the lower chamber and the plate was placed at 37°C in 5% CO_2_/air. The culture medium was changed every other day. ROCK inhibitor (10 µM, Y27632 STEMCELL#72304) was included in the culture medium for the first two days of culture. Organoids were counted and measured at day 14. Colony-forming efficiency (CFE) is calculated as the ratio between the numbers of spheres observed over the initial number (5000) of AT2 cells. At day 14, organoids were processed for whole-mount immunofluorescence staining.

### Whole-mount immunofluorescence staining of organoids

See supplementary methods

### Quantitative RT-PCR

See supplementary methods

### Co-staining: RNA in situ hybridization assay and IF

See supplementary methods

### Microarray analysis

See supplementary methods. GEO accession: <u>GSE162859</u>

### Statistics

All results are mean ± SEM. All error bars on graphs represent SEM. Statistical tests are 2-tailed *t* tests. *P* ≤ 0.05 was considered statistically significant.

### Patient and public involvement in research

n/a. This is a basic research project.

## Results

### The capacity of Cd45^Neg^Cd31^Neg^Epcam^Neg^ resident mesenchymal cells (rMCs) to functionally support the self-renewal and differentiation of alveolar epithelial type 2 (AT2) stem cells is associated with Stem cell antigen 1 (Sca1) expression

The aim of this study is to refine the adult lung resident mesenchymal cell population, called thereafter rMC and defined by flow cytometry as Cd45^Neg^Cd31^Neg^Epcam^Neg^ cells. This cell population is capable of sustaining AT2 stem cell renewal using the previously described alveolosphere in vitro assay [5]. In this approach, the use of antibodies against Cd45 and Cd31 allowed us to remove, from the target subpopulation, the hematopoietic and endothelial cells, respectively.

In the context of a previous milestone study generating organoids in Matrigel arising from a co-culture of Epcam^High^ Cd24^Low^ epithelial cells (previously called epithelial stem/progenitor cells or EpiSPC) with rMC, it was demonstrated that rMC expressing the stem cell antigen 1 (Sca1) were capable of sustaining the self-renewal and differentiation of these cells [13,14]. However, in this study, the impact of the different subpopulations of rMCs on AT2 stem cell self-renewal and differentiation was not analysed.

For this purpose, we have sorted an enriched population of mature AT2 cells from the *Sftpc*^*CreERT2/+*^; *Tomato*^*flox/+*^ lungs. We started first by isolating Cd31^Neg^Cd45^Neg^Epcam^Pos^ cells and then applied two successive and stringent sorting approaches: first using LysoTracker, a fluorescent dye that labels acidic compartments found abundantly in the lamellar bodies of AT2 cells and second, using the tomato reporter expressed specifically in AT2 cells (Fig. 1A). In addition, we sorted also rMCs-Sca1^Pos^ as well as rMCs-Sca1^Neg^ (Cd45^Neg^Cd31^Neg^Epcam^Neg^Sca1^Neg^). In order to have sufficient cells to carry out our assay, we pooled the lungs of four adult (6-8 weeks old) C57BL/6 mice. This allowed us to carry out 6 independent co-cultures. rMCs-Sca1^Pos^ represent around 25% of the total rMC population (Fig. 1A). When co-cultured with sorted mature AT2s, rMCs-Sca1^Pos^ led to the formation of organoids (Fig. 1B) with a colony forming efficiency of 5% (Fig. 1C) which is in line with previously published alveolosphere experiments. Interestingly, rMCs-Sca1^Neg^ fail to sustain alveolosphere formation (Fig. 1B). The difference in organoid size and CFE is highly significant between the two rMC subpopulations (Fig. 1C, *P* < 0.01).

**Figure 1:**
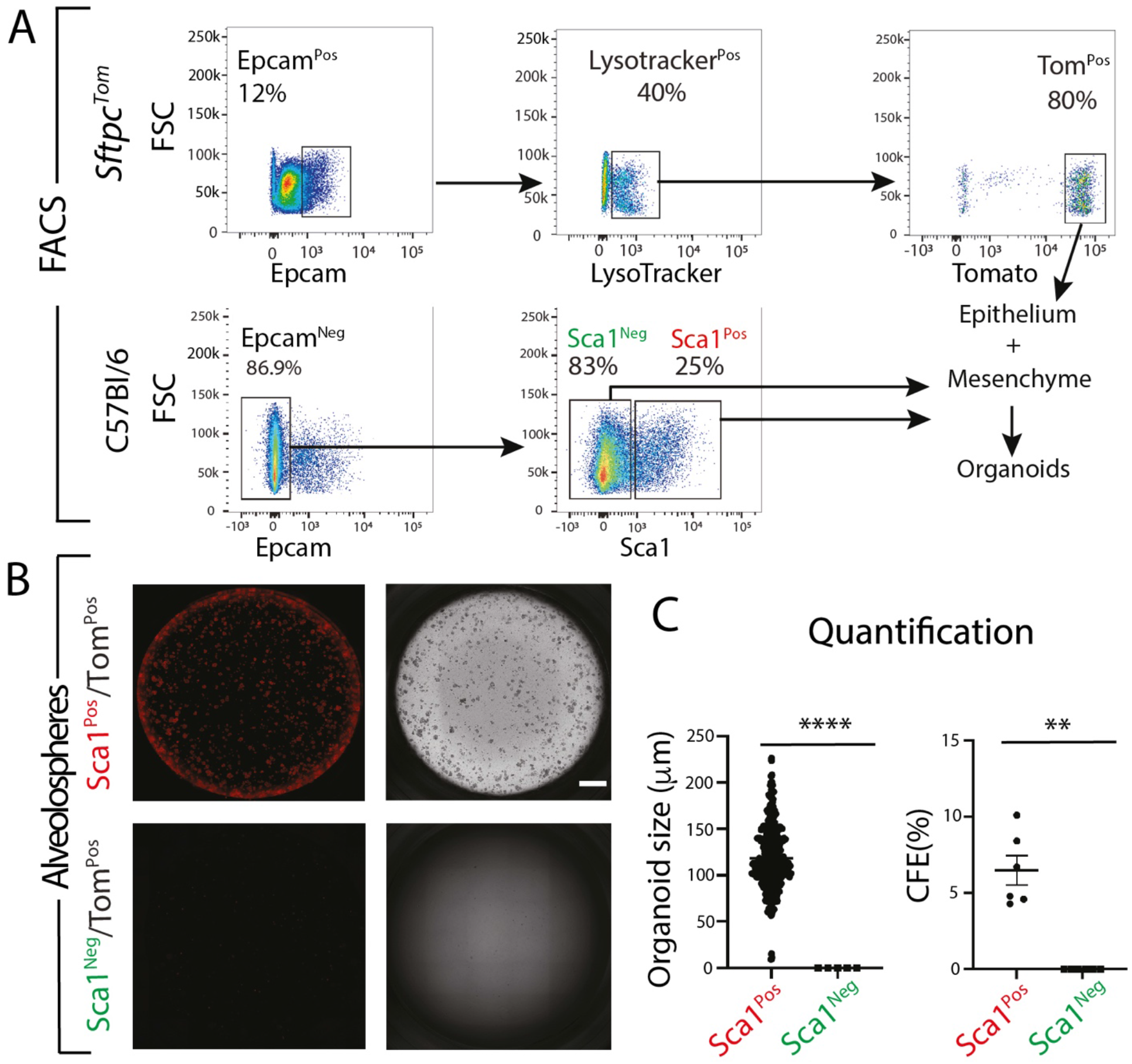
Sca1 expression in combination with the alveolosphere assay separates functionally the lung resident mesenchymal cells. **(A)** Gating strategy to sort mature AT2 (Epcam^Pos^Lysotracker^Pos^Tom^Pos^) from *Sftpc*^*Tom*^ as well as rMc-Sca1^Pos^ (Cd31^Neg^Cd45^Neg^Epcam^Neg^Sca1^Pos^) and rMC-Sca1^Neg^ (Cd31^Neg^Cd45^Neg^Epcam^Neg^Sca1^Neg^) cells from C57BL6 (WT) mice (**B)** Alveolosphere assay at day 14: Co-culture of mature AT2 with rMC-Sca1^Pos^ or rMC-Sca1^Neg^ cells (**C)** Quantification of the organoids; organoid size, and colony forming efficiency of six independent experiments (n=6). Scale bar; 50 µm

### rMCs-Sca1^Pos^ can be further functionally subdivided for the self-renewal and differentiation of AT2 stem cells on the basis of LipidTox staining

To better characterize rMCs-Sca1^Pos^ lung cells and in particular whether this population may include the lipofibroblasts (LIFs), we used LipidTOX (LT), an efficient fluorescent stain for neutral lipid. As LIFs are abundant in neutral lipid droplets, LT has been previously used to quantify LIF population during lung development [4].

Sorted rMCs-Sca1^Pos^LT^High^ and rMCs-Sca1^Pos^LT^Neg^ were tested functionally in the alveolosphere model for their capacity to induce the self-renewal and differentiation of AT2 stem cells (Fig. 2A). For this experiment, we pooled the cells from 3 mice and carried 4 independent co-cultures. When co-cultured with sorted mature AT2s, rMCs-Sca1^Pos^LT^High^ led to the formation of organoids (Fig. 2B) with a colony forming efficiency of 2% (Fig. 2C) while rMCs-Sca1^Pos^LT^Neg^ failed to sustain organoid formation. We also carried out at day 14, Immunofluorescence (IF) for LT (LIF marker), Hopx (AT1 cell marker) and DAPI in conjunction with the detection of the endogenous tomato reporter (Fig. 2B). Our results indicate LT^Pos^ cells are located in the periphery of each organoid. In addition, we observed abundant expression of Hopx within the organoid indicating proper AT2 to AT1 differentiation. RT-qPCR was also carried out to quantify the expression of *Fgf10* and *Pdfra*, two well-known markers enriched in LIFs (Fig. 2D). Our results indicate a trend towards an increase in the expression of *Fgf10* and *Pdgfra* in rMCs-Sca1^Pos^LT^High^ compared to rMCs-Sca1^Pos^LT^Neg^ (n=3 independent mice) (Fig. 2D). These results are in line with the previous observation that *Fgf10* and *Pdgfra* are also expressed in cells other than LIFs [4,15]. Overall, our data therefore support the previous conclusion that it is the LIF subpopulation of the rMCs that can preferentially support AT2 stem cell survival and differentiation.

**Figure 2:**
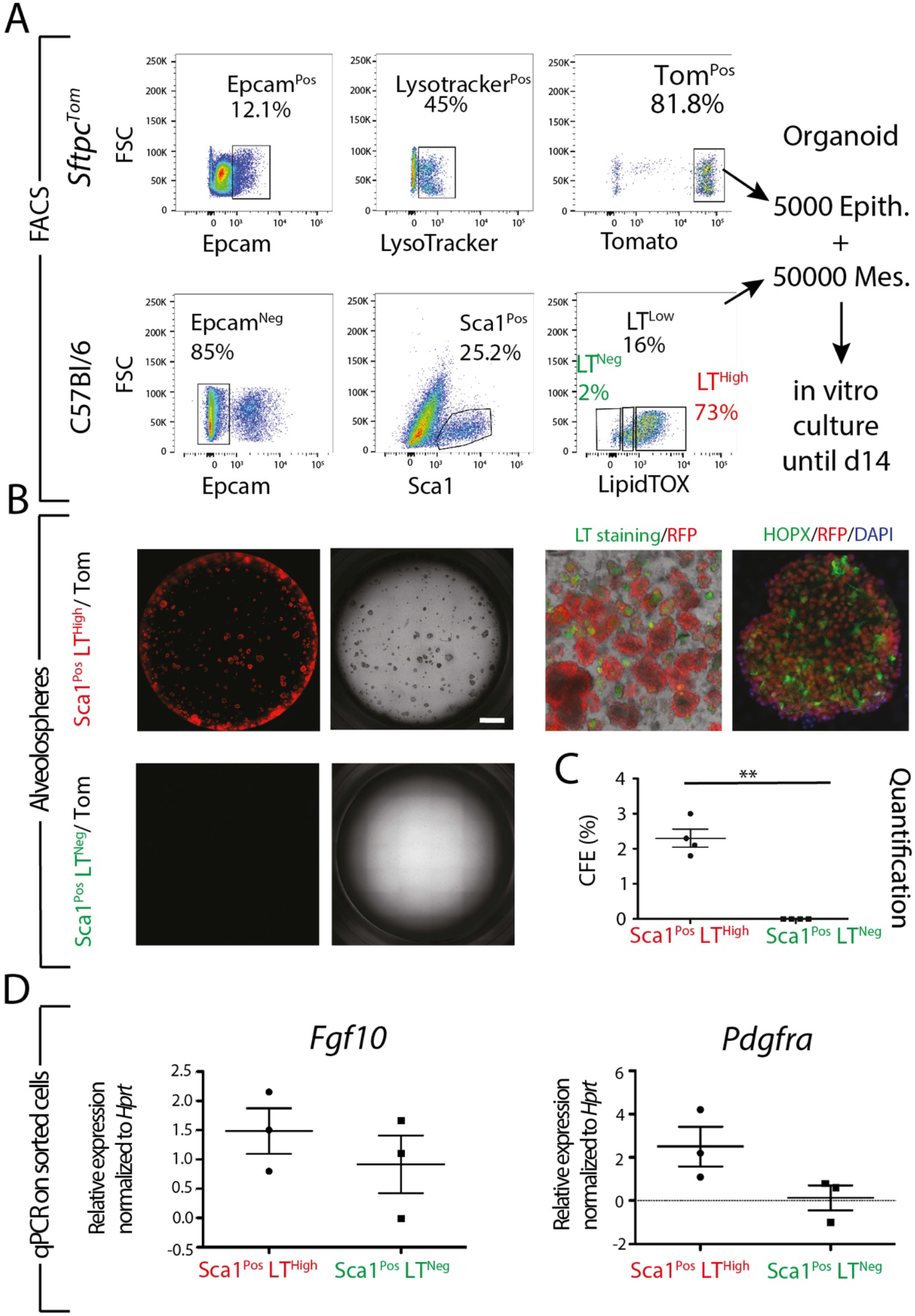
LipidTox staining identifies a subpopulation of rMC-Sca1^Pos^ cells supporting the self-renewal and differentiation of AT2 stem cells. **(A)** Sftpc^Tom^ and C57BL6 (WT) mice in 6-8 weeks of age were used to sort Lyso^Pos^Tom^Pos^ mature AT2 cells and rMC-Sca-1^Pos^LT^high^ or rMC-Sca-1^Pos^LT^Neg^ resident stromal cells. Mixture of cells were seeded in Matrigel in 24-well Transwell. **(B)** Fluorescence and bright field picture of representative wells at day 14. IF staining of organoids for DAPI, Sftpc (AT2 cell marker), and Hopx (AT1 cell marker) indicating that lipofibroblasts (rMC-Sca-1^Pos^LT^high^) support AT2 stem cell differentiation into AT1 cells (Scale bar 100 µm). **(C)** Quantification of colony forming efficiency (2.3 vs. 0 in rMC-Sca-1^Pos^LT^high^ vs. rMC-Sca-1^Pos^LT^Neg^, respectively (*P* value 0.001). **(D)** *Fgf10* and *Pdgfra* gene expression were assessed by qRT-PCR in rMC-Sca-1^Pos^LT^high^ vs. rMC-Sca-1^Pos^LT^Neg^. Scale bar; 50 µm

### Fgf10^Pos^ cells represent a niche for AT2 cells

We have previously reported that during development, a subset of Fgf10^Pos^ cells are progenitors for lipofibroblasts (LIFs) in the late stage of development and postnatally. We have also reported that during the early postnatal stage of lung development, only 28% of the LIFs express *Fgf10* indicating that the LIFs, like Fgf10^Pos^ cells [7], are an heterogenous population. To functionally evaluate the difference between rMCs-Sca1^Pos^Fgf10^Pos^ and rMCs-Sca1^Pos^Fgf10^Neg^ cells, we used the *Fgf10*^*LacZ*^ reporter line in combination with the ß-galactosidase fluorescent substrate FDG to sort, from rMCs, enriched populations of FDG^Pos^ (Fgf10^Pos^) and FDG^Neg^ (Fgf10^Neg^) cells. We then isolated the Sca1^Pos^ fraction for each subpopulation. Our results indicate that rMCs-Sca1^Pos^Fgf10^Pos^ represents around 33% of the rMCs-Fgf10^Pos^ and 25% of the rMCs-Sca1^Pos^. These results indicate that the rMCs-Sca1^Pos^ as well as the rMCs-Fgf10^Pos^ subpopulation are indeed heterogeneous (Fig. 3A). As previously described, we have co-cultured rMCs-Sca1^Pos^Fgf10^Pos^ as well as rMCs-Sca1^Pos^Fgf10^Neg^ cells with AT2 in our alveolosphere assay. Our results indicate that organoid only form with rMCs-Sca1^Pos^Fgf10^Pos^ with a CFE of 4% (n=4 independent experiments) (Fig. 3B and D). rMCs-Sca1^Pos^Fgf10^Neg^ are not capable of eliciting organoid formation. IF Staining of the organoids for Hopx show that the AT2 cells properly differentiate into AT1 cells (Fig. 3C).

**Figure 3:**
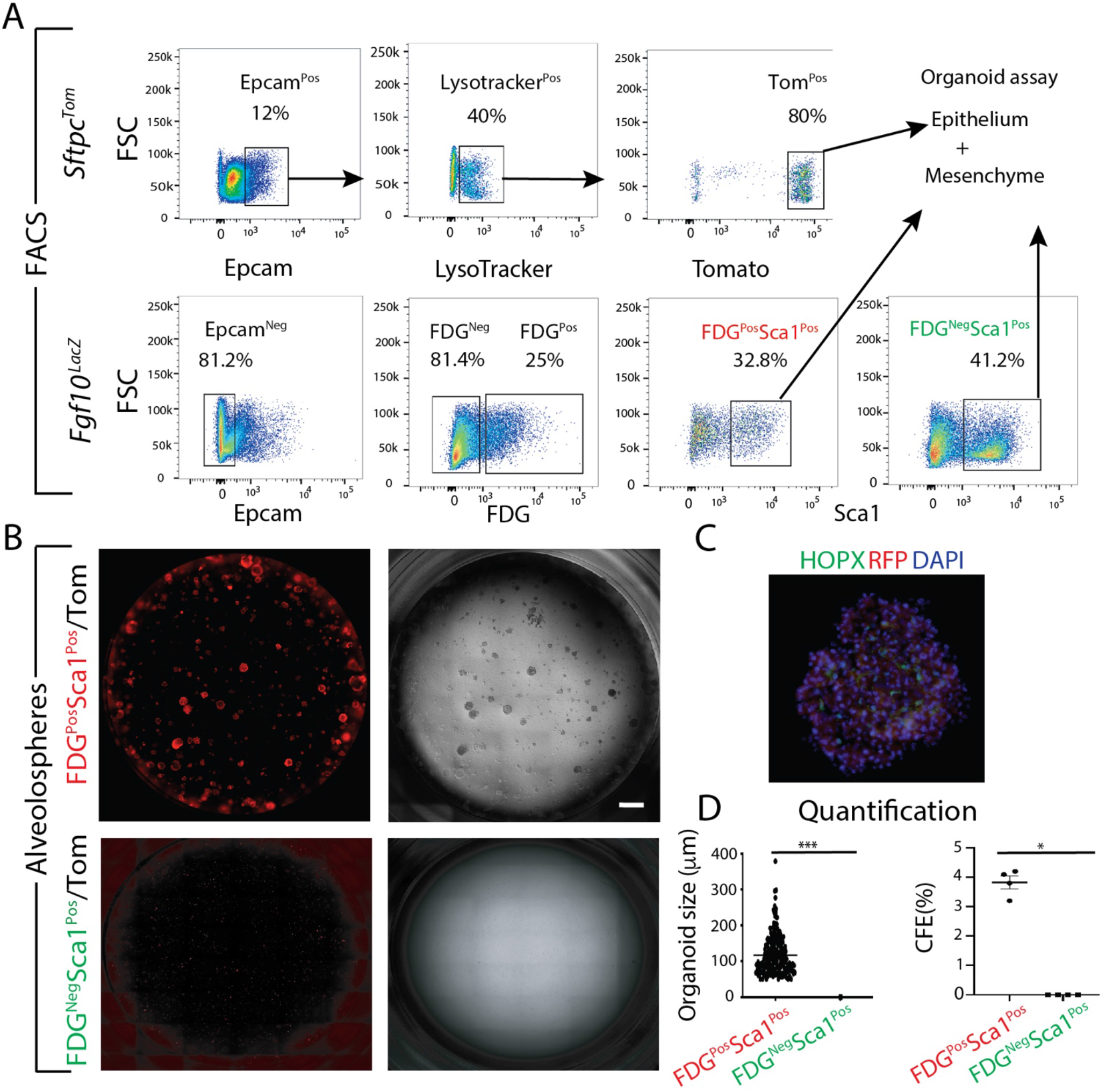
rMC-Sca1^Pos^Fgf10^Pos^ cells are supporting the self-renewal and differentiation of AT2 stem cells. **(A)** Gating strategy to sort mature AT2 cells from *Sftpc*^*Tom*^ lungs as well as rMC-Sca1^Pos^Fgf10^Pos^ and rMC-Sca1^Pos^Fgf10^Neg^ cells from *Fgf10*^*LacZ*^ lungs using the FDG fluorescent substrate for ß-galactosidase **(B)** Alveolosphere assay showing that only co-culture with rMC-Sca1^Pos^Fgf10^Pos^ leads to organoid formation. **(C)** IF staining against for Hopx and DAPI. **(D)** Quantification of organoid size and CFE with rMC-Sca1^Pos^Fgf10^Pos^ or rMC-Sca1^Pos^Fgf10^Neg^ (n=4). Scale bar;100 µm

### Comparison of rMCs-Sca1^Pos^Fgf10^Pos^ vs. rMCs-Sca1^Pos^Axin2^Pos^

Fgf10^Pos^ cells are progenitor for lipofibroblast. LIF in alveolar region are functionally important to support AT2 cells during lung development and postnatal stages. In addition, Axin2, a marker for Wnt signalling activation, is expressed in a subset of mesenchymal cells in the adult lung [16]. It has been reported that 74% of Axin2^Pos^ cells in the alveolar region are also expressing Pdgfrα. Axin2^Pos^ Pdgfrα^Pos^ mesenchymal cells are called mesenchymal alveolar niche cells (MANC) and are located closed to AT2 cells. Using the alveolosphere model they have been reported to sustain AT2 stem cell self-renewal and differentiation. As these cells have been described to express *Fgf7* and not *Fgf10* [8], we propose that rMCs-Sca1^Pos^Fgf10^Pos^ and rMCs-Sca1^Pos^Axin2^Pos^ represent two independent pools of niche cells for AT2 stem cells.

To better characterize these two rMC subpopulations, we used specific reporter lines; *Fgf10*^*LacZ*^ and *Axin2*^*LacZ*^ to monitor the distribution of LipidTOX staining in these two sub-lineages. By using FACSAria III cell sorter, we analysed 100,000 events, each sample contained harvested lung from one mouse -with the same age range (6-8 weeks old). Cd45^Neg^Cd31^Neg^Epcam^Neg^Sca1^Pos^ sorted cells were processed for further analysis (Fig. 4A). For *Fgf10*^*LacZ*^ lungs, we found 25% FDG^Pos^ (rMC-Sca1^Pos^Fgf10^Pos^) cells out of total rMC-Sca1^Pos^. For *Axin2*^*LacZ*^ lungs, our results indicate around 10% FDG^Pos^ (rMC-Sca1^Pos^Axin2^Pos^) cells out of total rMC-Sca1^Pos^. Further analysis based on LipidTox staining indicate that 85% of Fgf10^Pos^ and 98% of rMC-Sca1^Pos^Axin2^Pos^ cells were also LT^High^ cells (Fig 4B and C, respectively). Based on LT staining, we also report that most of the rMCs-Sca1^Pos^Fgf10^Neg^ as well as rMCs-Sca1^Pos^Axin2^Neg^ subpopulations contain a high percentile of LT^Low/High^ suggesting again a functional heterogeneity at the level of the LIFs (in regards to the maintenance of AT2 stem cell self-renewal) based on whether they express or not *Fgf10* or *Axin2*.

**Figure 4:**
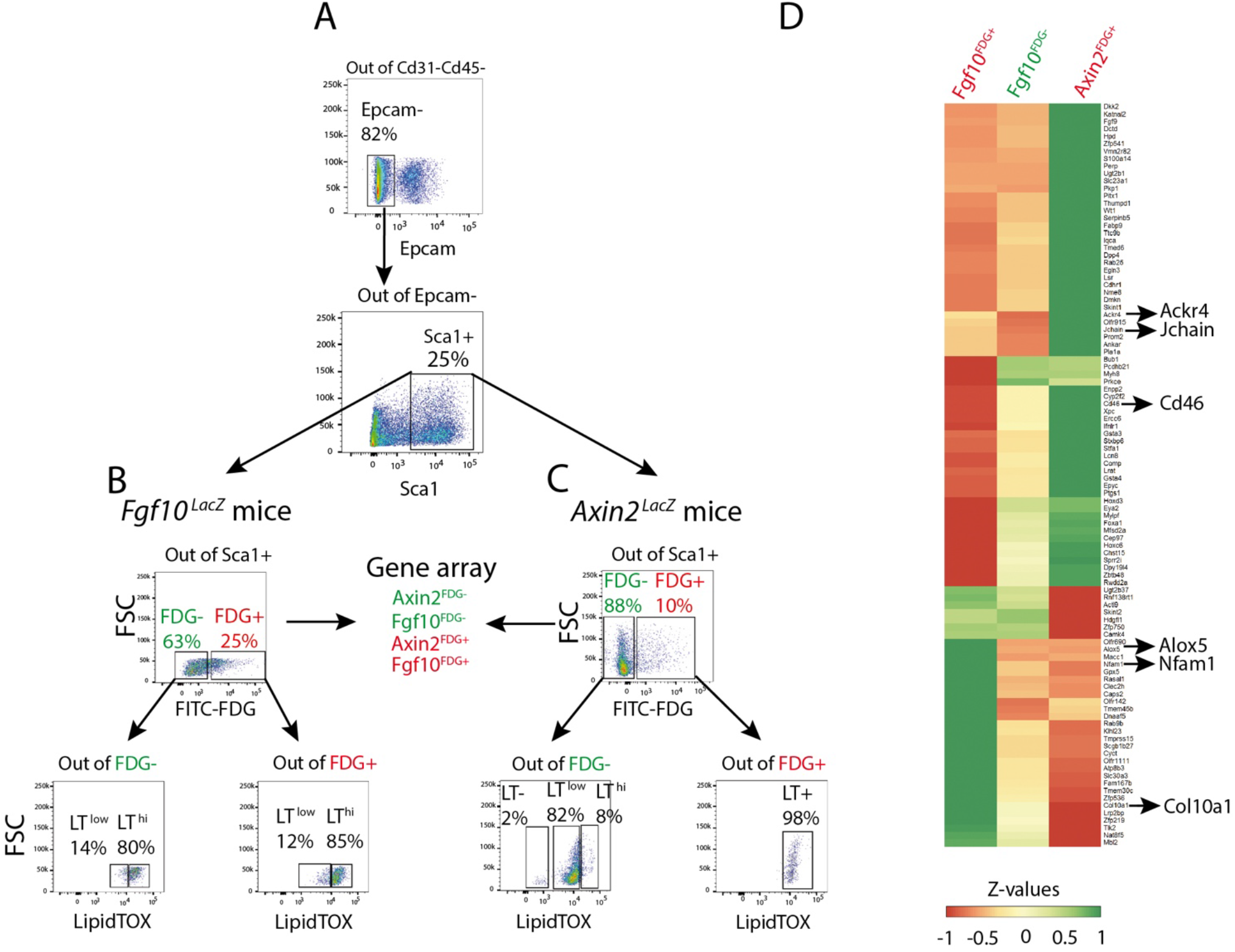
Comparison of rMC-Sca1^Pos^Fgf10^Pos^ vs. rMC-Sca1^Pos^Axin2^Pos^. **(A)** Single cell suspension from the adult lungs of *Fgf10*^*LacZ*^ *and Axin2*^*LacZ*^ reporter lines were processed by flow for Cd31, Cd45, Epcam and Sca1 expression to isolate rMC-Sca1^Pos^. (**B, C**) Staining with the FDG fluorescent substrate for ß-galactosidase allowed to separate further the rMC-Sca1^Pos^ into rMC-Sca1^Pos^Fgf10^Pos^ and rMC-Sca1^Pos^Fgf10^Neg^ (B) as well as rMC-Sca1^Pos^Axin2^Pos^ and rMC-Sca1^Pos^Axin2^Neg^ (C). Further staining with LipidTox allowed to quantify the abundance of LT^High^ in each subpopulation. **(D)** Heatmap is representing top 100 genes which are differentially expressed between rMC-Sca1^Pos^Fgf10^Pos^ and rMC-Sca1^Pos^Fgf10^Neg^. Note that the genes which are downregulated in rMC-Sca1^Pos^Fgf10^Pos^ are upregulated in rMC-Sca1^Pos^Axin2^Pos^ and vice versa. The arrows show some of the selected genes among the 100 genes.

In order to better define at the transcriptomic level the difference between rMCs-Sca1^Pos^Fgf10^Pos^ and rMCs-Sca1^Pos^Axin2^Pos^, we performed gene array analysis using the Agilent platform. Fig 4D shows top 100 genes which differentially expressed between rMCs-Sca1^Pos^Fgf10^Pos^ and rMCs-Sca1^Pos^Fgf10^Neg^ subsets. The genes differentially regulated between these two substets were then evaluated in rMCs-Sca1^Pos^Axin2^Pos^ cells. Our results indicate that several markers such as *Ackr4, Jchain, Cd46, Alox5, Nfam1* and *Col10a1* could be differentially expressed between rMCs-Sca1^Pos^Fgf10^Pos^ and rMCs-Sca1^Pos^Axin2^Pos^ and therefore used in the future to label these mesenchymal subpopulations. KEGG analysis of rMCs-Sca1^Pos^Axin2^Pos^ vs. rMCs-Sca1^Pos^Fgf10^Pos^ indicates an upregulation of metabolic pathways, RNA transport, DNA replication as well as cell cycle indicating that the Axin^Pos^ cells are metabolically active and proliferative (Fig. S1). Altogether, our data notably suggest that rMCs-Sca1^Pos^Fgf10^Pos^ cells are likely different from rMCs-Sca1^Pos^Axin2^Pos^ cells.

### *Fgf10* expressing cells are located close to Sftpc^Pos^ cells

To investigate the relative interaction between *Fgf10* expressing cells and AT2 cells, we combined the in-situ hybridization technique for detecting *Fgf10* mRNA expression with immunofluorescence staining for pro-SPC in adult wild type lungs. We found that <u>28%</u> ± 1% (n=3) of total *Fgf10* expressing cells are located close to pro-SPC expressing cells (Fig. 5A). This number is very similar to the one we reported before in the newborn lung using the *Fgf10*^*LacZ*^ reporter [4]. This observation suggests that a subset only of the *Fgf10* expressing cells constitutes a critical component of the alveolar niche that may robustly communicate with AT2 cells. To better define the heterogeneity of AT2 stromal niche, we have also performed co-staining of *Fgf7* mRNA and Sftpc. *Fgf7* expression has been proposed to be a hallmark of the MANC population. To validate the *Fgf7* mRNA probe, we used *Fgf7* KO adult lungs. While a clear signal was obtained with a positive control probe provided by the manufacturer, the *Fgf7* mRNA probe failed to generate a signal in the *Fgf7* KO lungs (Fig. S2) indicating that the signal observed in wild type lung is specific (Fig. 5B). We found that around only 2.0% ± 0.2% (n=3) of *Fgf7* expressing cells are in vicinity of Sftpc^Pos^ cells (Fig. 5B). Finally, we performed a co-staining for *Fgf7* and *Fgf10* mRNA using different fluorescent labelled probes. Our results indicate that 15.7% ± <u>3.0%</u> (n=3) of total cells are *Fgf7* expressing cells vs. <u>25.0%</u> ± <u>2.0% (n=3)</u> for *Fgf10* expressing cells. Finally, 4.0% ± <u>0.7% (n=3)</u> of total cells are double positive *Fgf10/Fgf7* cells (Fig. 5C). Taken together, these data suggest that the rMCs-Sca1^Pos^Fgf10^Pos^ are distinct from Fgf7^Pos^ MANC, while both subpopulations are located close to AT2 cells.

**Figure 5:**
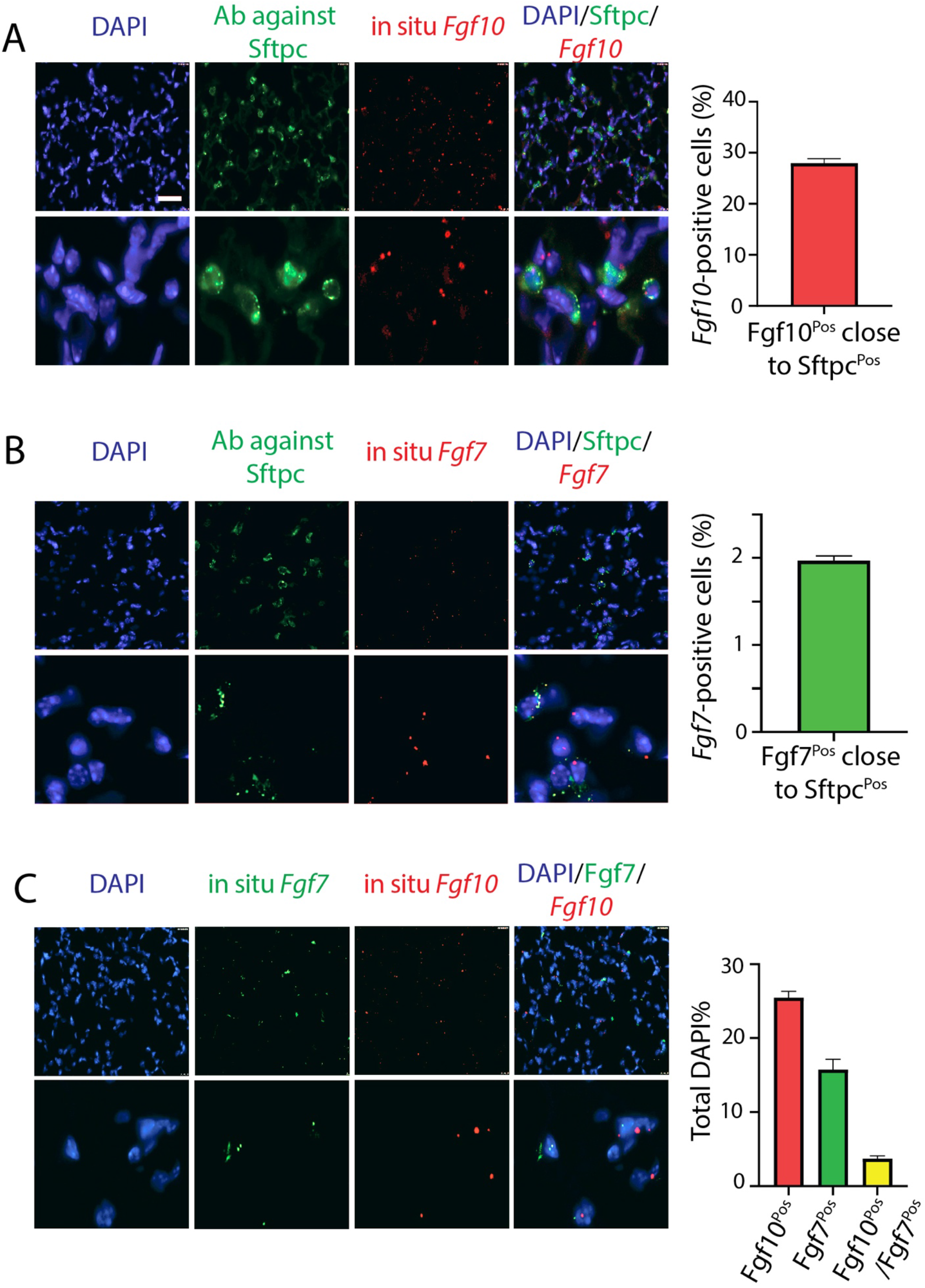
Analysis and comparison of *Fgf10* and *Fgf7* mRNA expression in relation to pro-Sftpc^Pos^ cells in the adult mouse lung. (**A**) In-situ hybridization for *Fgf10* mRNA (in red) and IF staining against for pro-Sftpc (in green). Low and high magnification. Quantification of *Fgf10* expressing cells close to Pro-Sftpc^Pos^ cells. **(B)** In-situ hybridization for *Fgf7* mRNA (in red) and IF staining against for pro-Sftpc (in green). Low and high magnification. Quantification of *Fgf7* expressing cells close to Pro-Sftpc^Pos^ cells. **(C)** Co-staining of *Fgf10* and *Fgf7* expressing cells. Quantification of *Fgf7, Fgf10* as well as *Fgf7/Fgf10* expressing cells compared to total cells. Scale bar for low magnification: 50 µm and Scale bar for High magnification 200 µm.

### rMCs-Sca1^Pos^ are impacted by obesity and gender

Patients who survived SARS-CoV-2 infection often develop lung fibrosis [9]. AT2 cells are proposed to be the main target of SARS-CoV-2 because of their widely distributed expression of angiotensin converting enzyme (ACE), a major receptor for this virus. As massive damages to the AT2s occur following the viral infection, it is likely that this process leads to the transdifferentiation of the LIFs into activated MYFs, thereby leading to fibrosis formation [1]. Obesity/diabetes and gender have been proposed to be aggravating pre-existing conditions predicting the severity of the disease [11].

To explore the impact of obesity/diabetes and gender on the functionality of the rMCs-Sca1^Pos^ cells, we used 6-8 weeks C57BL6 males and females as well as *Leptin*-deficient *Ob/Ob* (aka *Ob/Ob*) mutant male and female mice (n=2 for each gender and for WT and mutant). Fig. 6A shows the analysis by flow cytometry analysis of rMCs-Sca1^Pos^ in C57BL/6 mice vs. C57BL/6 Leptin-deficient *Ob/Ob* mice. Interestingly, a drastic reduction of the percentile of rMCs-Sca1^Pos^ is observed in *Ob/Ob* mice compared to wild type mice (27.7% vs. 9%, respectively). We also functionally tested the rMCs-Sca1^Pos^ from these different mice by co-culturing them with sorted AT2 cells using the alveolosphere assay. When the rMCs-Sca1^Pos^ are isolated from *Ob/Ob* male mice presenting the two risk factors, obesity and male (Fig. 6B), we observed a complete absence of organoid formation (n=2). However, when rMCs-Sca1^Pos^ are isolated from female *Ob/Ob* mice presenting only obesity as a risk factor, a significant number of organoids are present (3% CFE, n=2). This CFE is, however, lower than the one observed in non-obese C57BL6 female wild type mice (5% CFE, n=2) indicating that obesity alone is already impacting the functionality of the rMCs-Sca1^Pos^. Interestingly, non-obese C57BL6 male wild type mice display also a reduced CFE compared to non-obese C57BL6 female wild type mice (2% vs. 5%, respectively, n=2) indicating that the gender alone is also impacting the functionality of the rMCs-Sca1^Pos^. We also observed that age, another risk factor for COVID-19, was also having the same effect than obesity and gender on the capacity of the rMCs-Sca1^Pos^ to maintain the self-renewal of the AT2 stem cells (data not shown).

**Figure 6:**
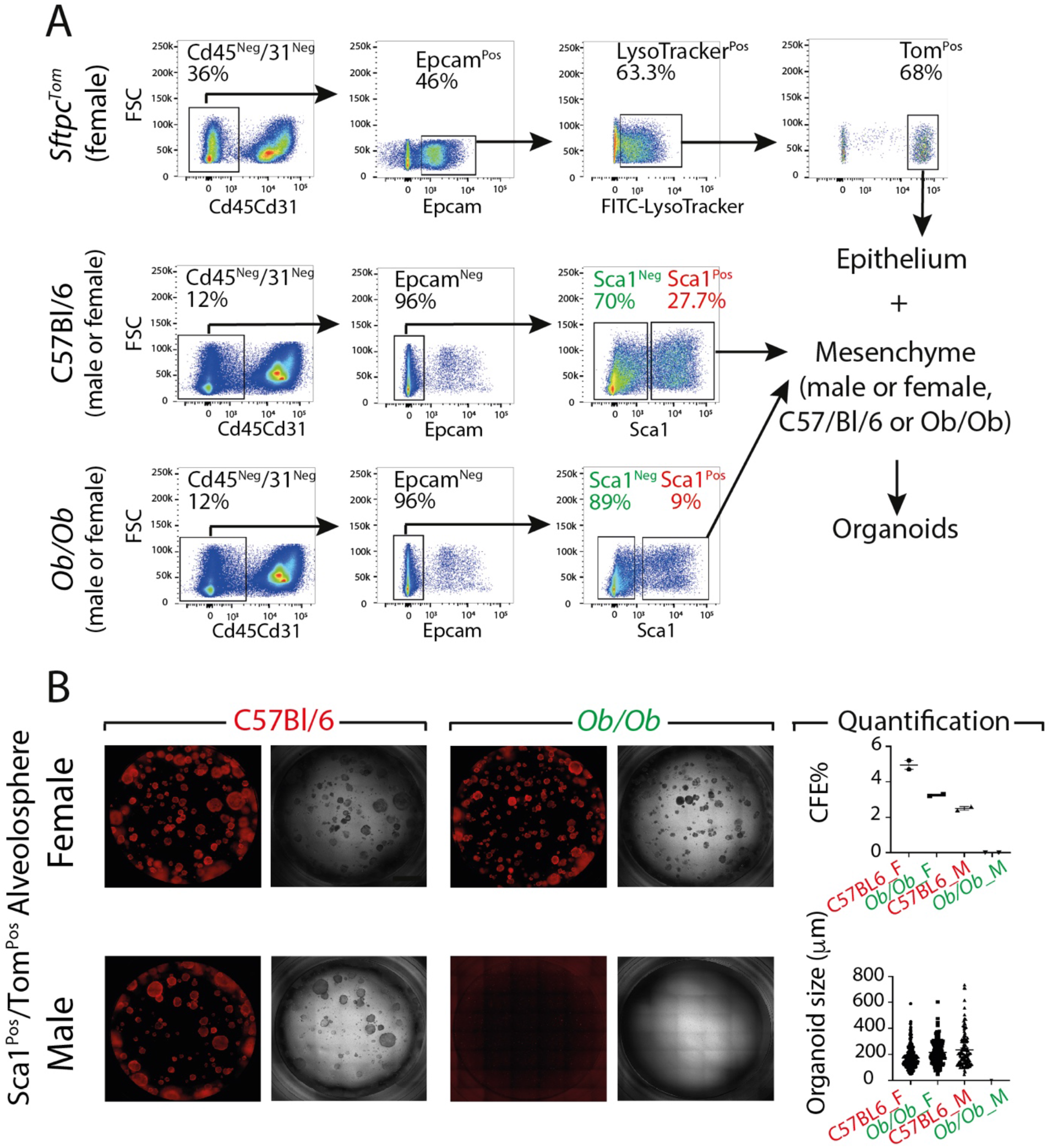
rMC-Sca1^Pos^ cells are affected by obesity and gender. (**A**) Flow cytometry analysis and gating strategy to sort mature AT2 cells from *Sftpc*^*Tom*^ as well as rMC-Sca1^pos^ from male and female C57BL/6 mice, and male and female *Ob/Ob* mice (n=2 for each genotype and gender). **(B)** Alveolosphere culture after 14 days and corresponding quantification of organoid size and colony forming efficiency (n=2 animals per condition). Scale bar; 50 µm

## Discussion

### rMC-Sca1^Pos^Fgf10^Pos^ are different from the rMC-Sca1^Pos^Axin2^Pos^

A major problem arising from key publications using flow cytometry as a primary experimental approach is the lack of independent studies reporting the reproducibility of the results. This is due in part to the type of flow sorter used, the sophisticated flow cytometry protocols, including the gating conditions used and the lack of comparison with other endogenous population at the time of the sort as internal controls. This is particularly crucial in the context of the lung mesenchyme which is still a huge black box both in human and in mice in spite of recent scRNA seq studies [17].In our study, we adopted an unbiased and reproducible FACS strategy to sort both epithelial and mesenchymal populations. We used Sca1 as a resident mesenchymal stem cell marker to refine the rMC population. In previous studies, Cd45^Neg^Cd31^Neg^Sca1^Pos^ cells (please note that Epcam was not used in that study) were detectable in the postnatal lung coincidently with the transition from the saccular to the alveolar stage of lung development. This population also co-expressed Cd34, Thy1 (Cd90), as well as Pdgfrα and was proposed to mark LIFs [18]. Interestingly, our gene array data shows that Cd90 is highly expressed in rMC-Sca1^Pos^Fgf10^Pos^ compared to rMC-Sca1^Pos^Axin2^Pos^ (data not shown). In another study, it was shown that Cd45^Neg^Cd31^Neg^Sca1^Pos^ could be refined into two main subpopulations, namely Cd166^Pos^Cd90^Neg^ and Cd166^Neg^Cd90^Pos^. The Cd166^Neg^Cd90^Pos^ subpopulation contained undifferentiated mesenchymal progenitors capable of differentiating towards the LIF and MYF lineage while the Cd166^Pos^Cd90^Neg^ subpopulation contained more differentiated mesenchymal cells already committed to the MYF lineage. Additionally, the Cd166^Neg^Cd90^Pos^ expressed high levels of *Fgf10* [14]. Taken together, our rMC-Sca1^Pos^Fgf10^Pos^ correspond to a specifically enriched population of LIFs capable of maintaining AT2 stem cell self-renewal and differentiation. A recent milestone study published in Cell described a mesenchymal cell subpopulation called MANC (Mesenchymal Alveolar Niche Cells), which was positive for *Axin2, Pdgfra, Wnt2, Il6* and *Fgf7* [8]. MANCs are located close to AT2s and sustain in vitro the self-renewal and differentiation of AT2 stem cells. The current knowledge, before this study was therefore that the MANC are considered at the top of a hierarchy of mesenchymal niche cells and are likely to be important for both homeostasis and repair after injury. The current study brings into light a novel challenger for this important role. Our results indicate that rMC-Sca1^Pos^Fgf10^Pos^ are likely different from the rMC-Sca1^Pos^Axin2^Pos^ (aka the MANC, but isolated in our experimental conditions) but has, nonetheless a similar function in regards to the AT2 stem cells.

### Are the rMC-Sca1^Pos^Fgf10^Pos^ more relevant than the rMC-Sca1^Pos^Axin2^Pos^ for the repair process after injury?

Given the fact that both subpopulations appear to sustain the self-renewal and differentiation of AT2 stem cells in vitro, a natural question is therefore whether they have redundant functions or whether one population appears to be more crucial than the other. From the angle of Fgf signalling and based on the consequence of *Fgf7* versus *Fgf10* gene inactivation in mice, we can conclude that rMC-Sca1^Pos^Fgf10^Pos^ are likely more important than rMC-Sca1^Pos^Axin2^Pos^. *Fgf10* inactivation leads to lung agenesis [19] while *Fgf7* null mice are viable and display no obvious lung phenotype [20]. Changes in endogenous *Fgf10* expression has been correlated with disease progression and/or repair after lung injury both in mice and humans [21–23]. Evidence for such a role for Fgf7 in the embryonic or adult lung in human or mice are still lacking in spite of the fact that Fgf7 was discovered 7 years before Fgf10 [24,25]. Another important question is whether it matters if the most efficient mesenchymal niche cells express Fgf7 versus Fgf10 as they are both ligands are acting through Fgfr2b.

Indeed, it does matter as Fgf7 and Fgf10, although belonging to the same Fgf subfamily of paracrine growth factors elicit different biological activities on isolated lung epithelium grown in Matrigel in vitro [6]. While Fgf10 induces the formation of new buds by a process of chemotaxis, Fgf7 triggers the proliferation of the epithelium leading to the formation of a cyst-like structure. Mass spectrometry studies demonstrated that Fgf10 differentially stimulated the phosphorylation of tyrosine 734 of Fgfr2b and the recruitment of the SH3-domain-binding protein 4 (SH3bp4) [26]. Tyrosine 734 phosphorylation controls the trafficking route of the receptor after internalization, allowing receptor recycling at the cell surface and sustained Akt and Shc phosphorylation. Knockdown of SH3bp4 or ectopic expression of a Y734F-mutated form of Fgfr2b in lung explants cultured in vitro modified the biological activity triggered by Fgf10 from chemotactic (bud formation) to proliferative (cyst-like structure).

### The activity of the rMc-Sca1^Pos^ cells is impacted by obesity and gender

In the context of lung fibrosis, the accumulation of activated MYF producing extracellular matrix components, modify the lung structure and negatively impact gas exchange. The LIF to MYF reversible differentiation switch appears to be a key process in fibrosis formation and resolution. Moreover, this transition was also shown *in vitro* in response to hyperoxia, as a key event in bronchopulmonary dysplasia (BPD) [27]. AT2 cells express angiotensin-converting enzyme II (ACE), a main receptor for SARS-CoV-2 [28]. Interestingly, SARS-CoV-2 induces the expression of transforming growth factor β (TGF-β) [29], which has been described to trigger the LIF to activated MYF transition. Such transition can be reversed by the administration of a PPARγ agonist (Rosiglitazone) as well as by the antidiabetic drug metformin [1,2]. Interestingly, metabolic dysregulation such as the one observed in obese patients has been associated with a worst prognostic in case of COVID-19. A similar conclusion has been reached for the gender as well as for the age. Interestingly, we can detect the impact of obesity, gender on the functionality of the rMC-Sca1^Pos^ cells to sustain AT2 stem cell self-renewal and differentiation. Our findings open the way for the future to screen for drugs capable of restoring the niche capabilities.

## Acknowledgment

We would like to thank Kerstin Goth and Jessica Nesswetha for the maintenance and management of the mouse colonies required for this study.

## Author contribution

ST has performed all experiments and MH has performed FACS analysis and sorting for samples. JW carried out the bioinformatics analysis. ST, JZ, SB, conceived the project and wrote the manuscript. All authors contributed to the article and approved the submitted version.

## Supplementary figures

**Figure S1:**
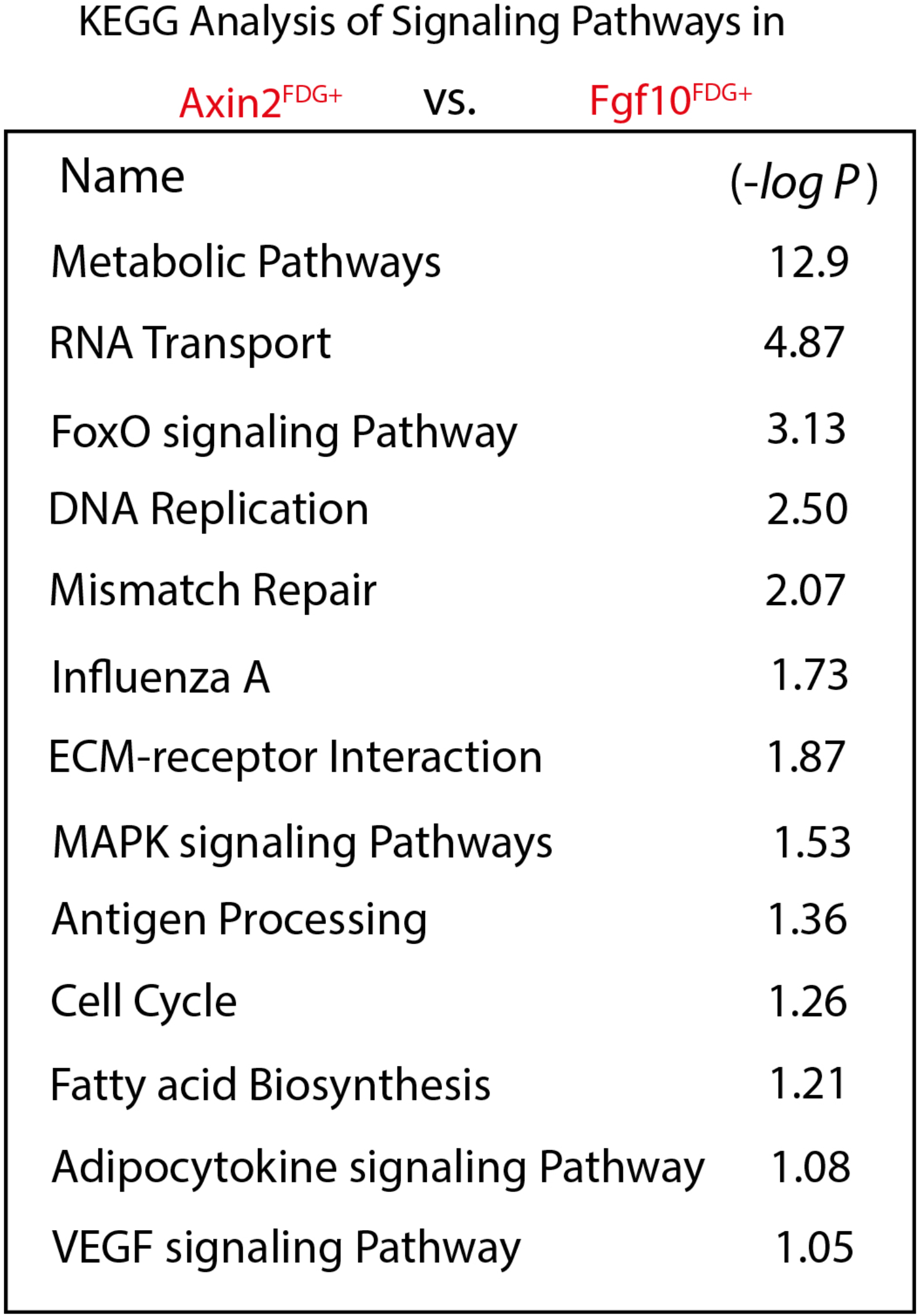
KEGG analysis of signalling pathways in Axin2^Pos^ cells vs. Fgf10^Pos^ cells.

**Figure S2:**
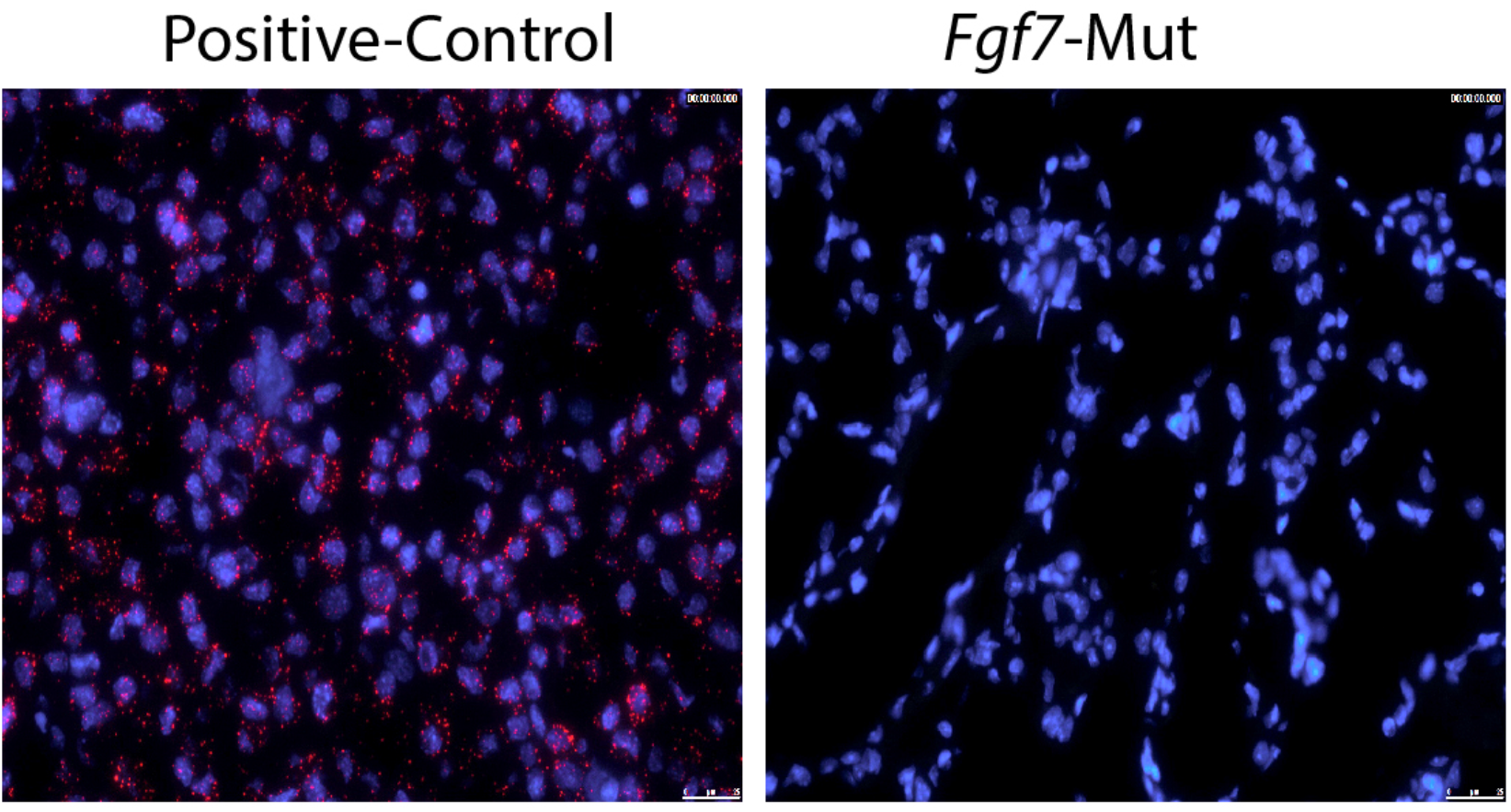
Validation of the *Fgf7* mRNA probe

